# Tau condensation on DNA and localization on centromeres: A potential link to cell division

**DOI:** 10.1101/2024.09.24.614852

**Authors:** Celine Park, Jaehun Jung, Yuri Hong, Chaelin Lee-Eom, Sang-Hyun Rah, Keunsang Yang, Jaehyeon Shin, Ayoung Jeong, Seokyun Hong, Jong-Bong Lee, Dong Soo Hwang, Min Ju Shon

**Affiliations:** Department of Physics, Pohang University of Science and Technology (POSTECH), Pohang, Republic of Korea; School of Interdisciplinary Bioscience and Bioengineering, Pohang University of Science and Technology (POSTECH), Pohang, Republic of Korea; Division of Environmental Science and Engineering, Pohang University of Science and Technology (POSTECH), Pohang, Republic of Korea

**Author notes:** These authors contributed equally to this work (Celine Park, Jaehun Jung). Present address: Max Planck Institute of Cell Biology and Genetics, 01307 Dresden, Germany (Yuri Hong).

## Abstract

Tau protein, traditionally recognized for stabilizing microtubules and forming pathological aggregates, has recently been observed to form condensates in various contexts. While its condensation with RNA has been well studied, the interaction between tau and DNA, along with its biological significance, remains less explored. Here, using single-molecule experiments, we found that tau binds stably to naked DNA at nanomolar concentrations, leading to the local co- condensation of tau and DNA. These tau condensates on DNA can also interface with microtubules, leveraging tau’s known role in promoting microtubule growth and organization. The dynamic nature of these condensates facilitates the remodeling of the DNA–microtubule assembly. Interestingly, two phosphomimetic tau mutants, T231D/S235D and S262D, retained their affinity for DNA but differed in their ability to link microtubules to DNA. Furthermore, imaging of HEK-293 and SH-SY5Y cells in early mitosis revealed that tau localizes on centromeres, poised to capture nascent mitotic spindles. Building on these observations, we speculate that tau may play a novel role in mitosis, where tau clusters facilitate the early registration of mitotic spindles to chromosomes before kinetochore-mediated attachment. We also discuss the potential implications of this model in conditions where abnormal cell cycle re-entry and tau activity may disrupt cell division.

## Introduction

Tau protein has long been known for its primary role in stabilizing microtubules (*1*, *2*). More recently, its ability to facilitate the nucleation and growth of microtubules has gained attention (*3*). This function relies on tau’s tendency to form condensates at high concentrations (*4–6*), a mechanism increasingly recognized as an organizing principle in many areas of cell biology (*7*, *8*). At the same time, tau has been a major focus of research in neurodegenerative diseases due to its critical involvement in forming pathological aggregates (*9*). The abnormal aggregation of tau is a hallmark of tauopathies, including Alzheimer’s disease, and is strongly associated with neuronal dysfunction and cell death.

In addition to binding microtubules, tau’s multifaceted biophysical properties enable a variety of dynamic interactions (*10*). As an intrinsically disordered protein with distinct domains (*11*), tau adopts flexible conformations, allowing for weak multivalent interactions with both itself and other molecules. This ability is crucial for driving the co-condensation of other factors within tau condensates. One example is tubulin, which has been shown to be recruited and to locally nucleate microtubule growth (*3*). Another is the complexation between tau and RNA, which has been explored in the context of electrostatic interactions (*4*, *12*), participation in stress granules (*13*), and the development of tau fibrils and aggregates (*5*, *14*, *15*). As a corollary, the interactions between tau, tubulin, and RNA can also modulate tau’s ability to organize microtubules (*16*). These examples suggest that tau condensates may act as hubs for various biomolecules sparking distinct cellular activities.

Unlike tau–RNA interactions, the interaction between tau and DNA has received less attention. Since tau primarily resides in the cytoplasm, its limited access to nuclear DNA may have reduced the perceived physiological relevance of tau–DNA binding. Nevertheless, interactions between tau and naked DNA molecules have been reported (*17–23*), albeit infrequently. While it is unsurprising that tau may bind to dsDNA through a mechanism similar to its interaction with RNA, the exact nature of tau–DNA interactions likely differs due to the intrinsic stiffness of dsDNA. The biophysical strength of this interaction, as well as its potential role in living cells, remains largely unexplored. One notable exception is tau’s role in the nucleus, where it has been proposed to protect the genome and influence gene expression (*24–26*).

Here, we report that tau directly interacts with DNA, forming co-condensates that serve as dynamic platforms for microtubule organization. Our single-molecule observations reveal that 2N4R tau, the longest isoform of tau expressed in the adult human brain, forms mobile condensates on DNA strands at low concentrations, which subsequently bind to microtubules. These tau–DNA co-condensates act as dynamic microtubule attachment sites, assembling microtubules around the DNA, with phosphomimetic mutations (T231D/S235D and S262D) adding further complexity to this process. Furthermore, by imaging tau-expressing cells during division, we observed tau clustering around the centromeres of mitotic chromosomes during prometaphase, poised to interface with nascent mitotic spindles. Based on these findings, we speculate a new role for tau in cell division, extending beyond its nuclear function. We hypothesize that as the nuclear envelope breaks down and chromosomes become exposed to cytoplasmic tau, the dynamic rearrangement of tau on mitotic chromosomes facilitates the “search and capture” process of newly forming microtubules—a process traditionally considered random. This model further proposes that as tau–DNA co-condensates mature and migrate toward the centromeres, these transient attachment sites are gradually replaced by kinetochores for a more stable connection.

## Results

### Tau stably binds to naked DNA at nanomolar concentrations

To probe the interaction between tau protein (2N4R, tau-441) and naked DNA, we first examined the mobility of DNA in an agarose gel after mixing it with purified tau protein (Fig. S1). The prepared dsDNA samples of varying lengths (24–5,000 bp) all displayed a significant mobility shift when incubated with wild-type tau (500 nM or higher) (Fig. S2), indicating a strong association between tau and DNA. This interaction was further enhanced when the NaCl concentration was lowered from 100 mM to 25 mM, suggesting an electrostatic nature of the binding, similar to tau– RNA complexation (*4*). Given that the shortest DNA we prepared (24 bp) allowed tau binding, it is likely that the longer DNA constructs can accommodate proportionally more tau molecules along their length.

While this bulk assay confirmed tau’s DNA-binding properties, it did not reveal the distribution of tau within and across DNA strands, as well as the impact of this binding on DNA conformation. To address these questions, we set up a series of single-molecule imaging experiments that directly visualize the binding of tau protein to DNA substrates. We first performed a classic flow-stretching experiment in a flow chamber, where 48.5-kbp λ-DNA molecules were attached to a polyethylene glycol (PEG)-coated glass coverslip by one end and then stretched upon buffer flow (7 μL/min) (Fig. 1A). The stretched DNA strands were visualized using a fluorescent stain (SYTOX Orange) on a total internal reflection fluorescence (TIRF) microscope (Fig. 1B). The tethered λ-DNA molecules were gently stretched to 6.2 ± 0.6 μm (∼40% of the contour length) (Fig. 1C, D), indicating ∼0.1 pN of flow-induced stretching force. When 500 nM tau (10% labeled with Cy5) was added at the same flow rate, the λ-DNA molecules gradually shortened and collapsed onto the attachment point (Fig. 1B–D and Movie S1). We checked that the fluorescently labeled tau also accumulated on the condensed DNA spots (Fig. 1B, E), indicating that tau binding is the direct cause of DNA compaction. Note that these observations were made without highly concentrated tau solutions or a crowding agent, both of which are typically needed to induce condensation behavior through liquid-liquid phase separation (LLPS) (*5*, *6*, *27*). This indicates that a stable interaction between tau and naked DNA, which locally enriches tau, is sufficient for the observed DNA compaction.

**Fig. 1.**
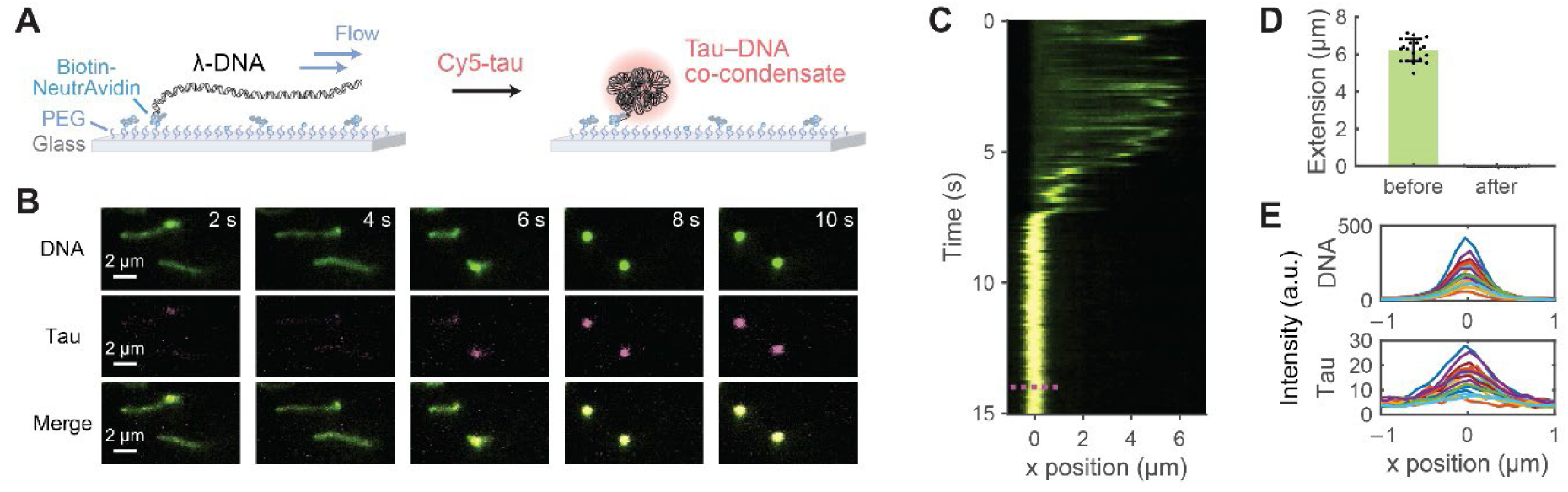
Condensation of DNA by tau (A) Flow stretching of surface-tethered λ-DNA, followed by the formation of tau–DNA co-condensates upon the addition of Cy5-labeled tau. (B) Time-lapse images from TIRF microscopy showing the co-condensation event during the addition of 500 nM tau. DNA is stained with SYTOX Orange (*green*), and tau is labeled with Cy5 (*magenta*). (C) Representative kymograph of the co-condensation event. The magenta dotted line indicates the region from which the fluorescence intensity profiles were sampled, as shown in (E). (D) Extension of flow-stretched λ-DNA before and after tau addition (*n* = 20 molecules; mean ± s.d.). (E) Fluorescence intensity profiles of DNA (*top*) and tau (*bottom*) after tau addition, sampled from multiple locations along the DNA and aligned to their respective attachment points (*n* = 20 molecules). The colocalization of tau and DNA fluorescence indicates the formation of co-condensates.

### Tau binding to DNA generates local co-condensates

Recent discoveries indicate that DNA-binding proteins can form condensates on DNA, thereby compacting and remodeling its underlying structure (*28–34*). Motivated by these findings, we hypothesized that tau could exert similar effects on DNA. To test this, we prepared λ-DNA molecules tethered at both ends to a surface and introduced Cy5-labeled tau again (Fig. 2A), simultaneously imaging the fluorescence of both tau and DNA. As observed in single-tether experiments, there was significant fluorescence from tau binding to the DNA (Fig. 2B, C, and Movie S2). The DNA backbone was diffusely coated with tau, interspersed with multiple bright foci. As expected from the bulk gel shift assay, tau binding was both tau and salt concentration-dependent (Fig. 2C). Higher concentrations of tau resulted in more intense spots, even though we kept the concentration of the Cy5-labeled tau constant (50 nM) across all samples.

**Fig. 2.**
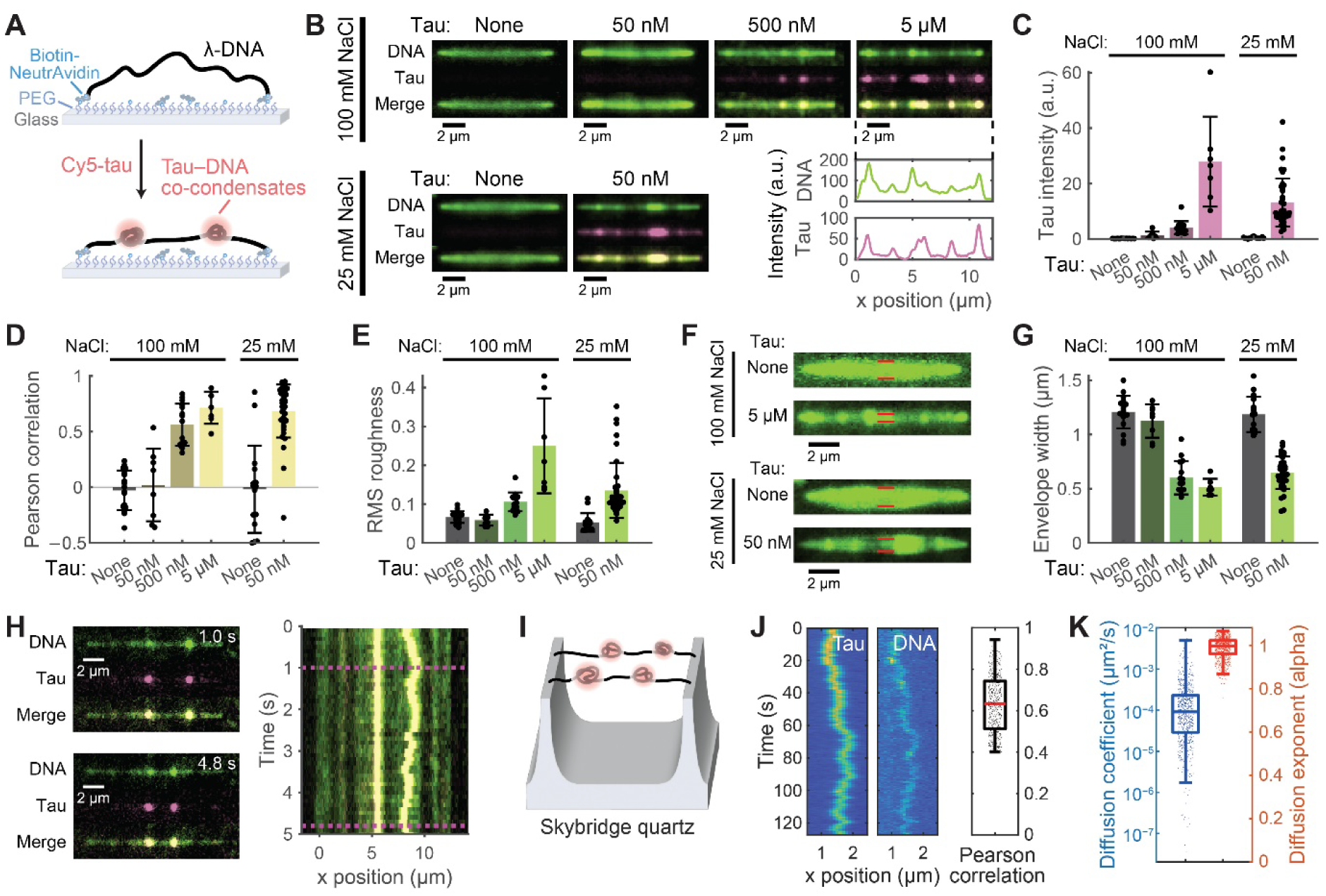
Tau–DNA co-condensates on DNA strands (A) Schematic of tau–DNA co-condensation on doubly tethered λ-DNA. (B) TIRF images of λ-DNA with different tau concentrations at 100 mM and 25 mM NaCl. DNA is labeled with SYTOX Orange (*green*), and tau with Cy5 (*magenta*). Intensity profiles for 5 μM tau (*bottom right*) high correlation between DNA and tau. (C) Quantification of tau fluorescence intensity for the conditions shown in (B). (D) Pearson correlation coefficients between DNA and tau fluorescence, indicating colocalization. (E) Root mean square (RMS) roughness of DNA fluorescence, reflecting puncta formation upon tau binding. (F) Maximum intensity projections of DNA fluctuations (over 5 s), showing the effect of tau on DNA flexibility. Red lines indicate the measured envelope width. (G) Distribution of DNA envelope width as a function of tau and NaCl concentrations, reflecting changes in DNA tension. (H) Snapshots and kymograph of mobile tau–DNA co-condensates. Magenta dotted lines indicate snapshot locations. These images show that tau and DNA move together, indicating the mobility of the co-condensates. (I) Schematic of the DNA skybridge technique for visualizing condensate mobility. (J) Kymograph of tau and DNA movement, with a corresponding Pearson correlation plot. (K) Box plots of diffusion coefficients and exponents of tau–DNA co-condensates, showing their diffusive behavior. Data in (C)–(G) are from *n* = 7 or more molecules for each condition (mean ± s.d.), and data in (J)–(K) are from *n* = 623 spot; box plots show the median (central line) and interquartile range, with whiskers extending to the minimum and maximum values not considered outliers.

Although the DNA coverage by tau varied among DNA strands due to different levels of foci formation, the locations of these bright regions corresponded closely between DNA and tau signals (Fig. 2B inset and Fig. 2D), suggesting that clustered tau binding leads to local compaction of DNA. The level of local compaction was estimated by measuring the RMS roughness of the DNA fluorescence signal along the strands, which showed a gradual increase as more tau was added (Fig. 2E). In contrast, DNA strands without tau showed uniform fluorescence along their length. Consistent with the local compaction effect, the binding of tau noticeably reduced the thermal fluctuation of DNA (Fig. 2F and Movie S2), indicating that the DNA became shorter and tauter. The mean envelope width, or the amplitude of off-axis deviation due to fluctuation, gradually decreased as tau concentration increased from 50 nM to 5 μM (Fig. 2G), consistent with the increase in tau binding. Notably, even 50 nM tau was sufficient to induce strong condensation of DNA under low-salt conditions (25 mM NaCl), as demonstrated by both foci formation (Fig. 2B, E) and reduced fluctuation (Fig. 2G). These results demonstrate that wild-type tau protein at low concentrations can stably bind to naked DNA molecules under physiological conditions.

To ensure that our findings on tau–DNA co-condensation extend beyond single-molecule observations, we checked bulk co-condensation between high concentrations of tau and dsDNA in the presence of 10% PEG, a standard method to simulate molecular crowding and drive phase separation. Under these conditions, tau and DNA readily formed co-condensates across various concentrations of tau, DNA, and salt (Fig. S3), consistent with a recent report on this aspect (*35*). Importantly, the co-condensation behavior was robust across a wide range of DNA lengths, from 24 bp to λ-DNA, indicating that DNA length did not significantly influence its partitioning into tau droplets. This tau-driven DNA condensation, reversing the above single-molecule observations of DNA-driven tau condensation, reinforces the strong molecular interaction between tau and DNA.

### Tau–DNA co-condensates are mobile

Surprisingly, despite the strong affinity between tau and DNA, many of the tau condensates remained mobile along the DNA substrates (Fig. 2H and Movie S3). Moreover, the DNA foci formed by tau also diffused together, irrespective of the tau concentrations used (50 nM – 5 μM). This suggests that, in addition to the tightly bound tau-DNA complexes within the core, a more loosely associated fraction of tau remains dynamic and capable of modulating DNA architecture through rearrangement, reminiscent of the liquid-like behavior characteristic of LLPS-driven condensates. However, not all condensates exhibited uniform mobility, likely due to heterogeneity in their mechanical properties (*e.g.*, differential maturation to more solid-like states), friction from contact with the PEG-coated glass surface, or a combination of both.

To alleviate the surface-imposed friction, we re-evaluated the mobility of tau–DNA condensates on the DNA skybridge platform (Fig. 2I) (*36*). In this technique, λ-DNA is suspended between microfabricated quartz barriers, reducing surface interaction artifacts and enabling high-throughput visualization of condensate mobility. Indeed, by allowing sufficient spacing between the DNA strands and the substrate surface, we confirmed that the tau and DNA foci migrated together along the DNA strands (Fig. 2J and Movie S4). Essentially all of the observed co-condensates (*n* = 623 spots) exhibited normal diffusion (diffusion exponent of 1) with large diffusion coefficients of 10^(−4.1^ ^± 0.7)^ μm^2^/s (Fig. 2K). In conclusion, these findings indicate that tau binding to DNA leads to dynamic co-condensation, which spatiotemporally modulates the local compaction of DNA.

### Tau binding exerts piconewton condensation force on DNA

The flow-stretching and double-tether experiments showed that tau binding can induce DNA condensation, counteracting a force of at least ∼0.1 pN from the buffer flow. We next employed magnetic tweezers to accurately measure the force generated by tau condensates on DNA and to determine if the condensation can be mechanically reversed (Fig. 3A). A 10-kbp dsDNA fragment was prepared as a substrate, with one end attached to a PEG-coated glass surface and the other to a magnetic bead for manipulation. Changes in DNA extension were monitored by tracking the vertical movements of the bead. This setup was also equipped with a TIRF microscopy module (*37*), which checked the fluorescence signal from tau molecules binding to DNA (Fig. S4).

**Fig. 3.**
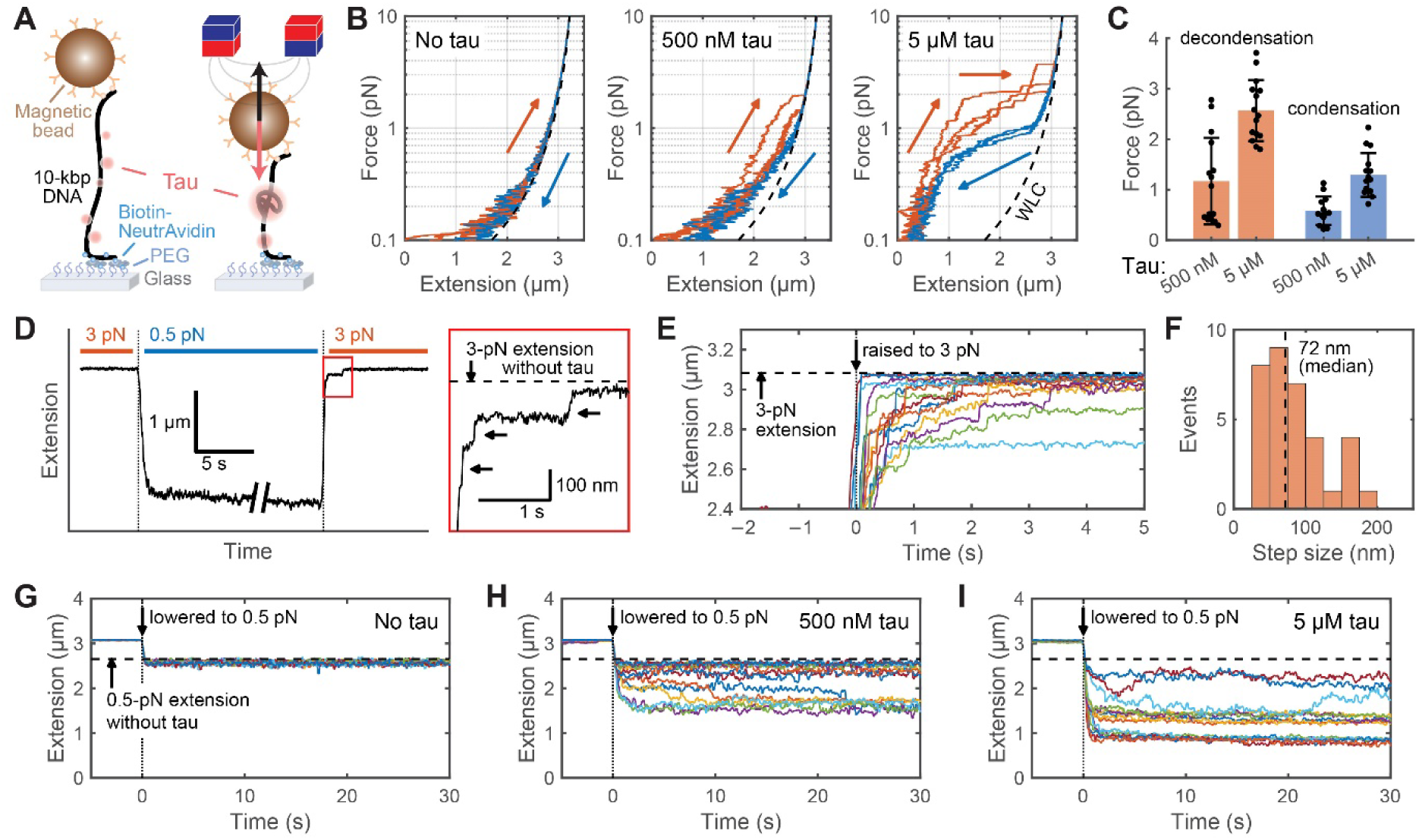
Mechanical characterization of tau–DNA co-condensates using magnetic tweezers (A) Schematic of magnetic tweezers experiments to probe tau–DNA co-condensation. (B) Representative force-extension curves of DNA during stretching (*orange*) and relaxation (*blue*) in the absence and presence of tau (*n* = 3 cycles each). Dashed line shows the worm-like chain model for dsDNA. (C) Condensation and decondensation forces obtained from force-extension measurements (*n* = 15 events; mean ± s.d.), as shown in (B). (D) Force-jump trace for 5 μM tau, showing compaction at 0.5 pN and stretching at 3 pN. The region in the red box is expanded on the right to highlight multiple steps observed during the 3 pN application. (E) Extension time series for 5 μM tau at 3 pN (*n* = 15 trials), aligned at the force jump. (F) Step size histogram from (E) (*n* = 34 events). (G–I) Extension time series at 0.5 pN for different tau concentrations (*n* = 15 trials each).

In the absence of tau, the classic worm-like-chain extension of dsDNA was reproduced by applying a force ramp between 0.1 and 10 pN (Fig. 3B, *left*). To ensure that the DNA constructs were free of single-stranded nicks, we used only those that collapsed under applied torque, exhibiting supercoiling behavior (Fig. S4). However, when 500 nM tau was added to the buffer, the overall extension of DNA was notably reduced, especially in the low-force range between 0.1 and 3 pN (Fig. 3B, *center*), consistent with tau-induced condensation of DNA. Fluorescence verification confirmed the accumulation of tau molecules on the DNA strands (Fig. S4, Movie S5). At higher forces, in contrast, the extension returned fully to that of naked dsDNA, implying that the tau molecules were largely displaced by stretching the DNA. It was possible to drive this compaction and decompaction cycle by repeatedly stretching and relaxing the DNA. Although the exact force levels at which these transitions occurred varied across trials (Fig. 3C), slightly higher forces (1–3 pN) were required than those at which condensation first occurred (<1 pN), suggesting some mechanical hysteresis and energy dissipation in the tau–DNA interactions. Finally, increasing the concentration of tau (from 500 nM to 5 μM) raised the condensation force, consistent with an increased tendency to undergo condensation.

The force-extension curves in the presence of tau revealed both smooth and stepwise changes in DNA extension. We considered whether the stepwise changes might represent a specific unit length of DNA compacted by tau condensates of relatively uniform size. To better resolve these events, we applied force-jump cycles (Fig. 3D), where the force was abruptly switched between 0.5 and 3 pN to trigger compaction and decompaction events. During the stretching phase, we detected many stepwise length changes (Fig. 3E), with a median size of 72 nm (Fig. 3F). In contrast, during the condensation measurements at 0.5 pN, smooth, gradual compaction dominated, with extension fluctuations over a longer period (∼1 min), indicating a weaker, more dynamic association phase.

### Tau condensates facilitate microtubule attachment to DNA

The tau condensates formed on DNA substrates are reminiscent of those that facilitate microtubule growth and assembly (*3*). Presumably, the DNA-bound tau condensates might function similarly to organize microtubules, possibly by cooperating with tau molecules already present on the microtubule surface. To test this idea, we introduced a mixture of tau and tubulin over the surface-bound tau–DNA assembly and evaluated whether the tau condensates maintain their potential to organize a microtubule network. We first checked that mixing tubulin with tau in the presence of GTP and 5% PEG induced rapid polymerization of microtubules (Fig. S5). When this solution of nascent microtubules was introduced over surface-tethered DNA molecules prepared at a low density, tau–DNA co-condensates effectively captured them (Fig. 4), suggesting that tau retains its essential microtubule binding activity. Importantly, fluorescence from free tubulin also localized within the tau condensates (Fig. 4A), indicating that these spots might serve as seeds for nucleating new microtubules. The microtubules engaged with the DNA strands either along their sides or via their ends, analogous to the lateral and end-on attachments posited for mitotic spindle coupling to chromosomes during mitosis (*38*).

**Fig. 4.**
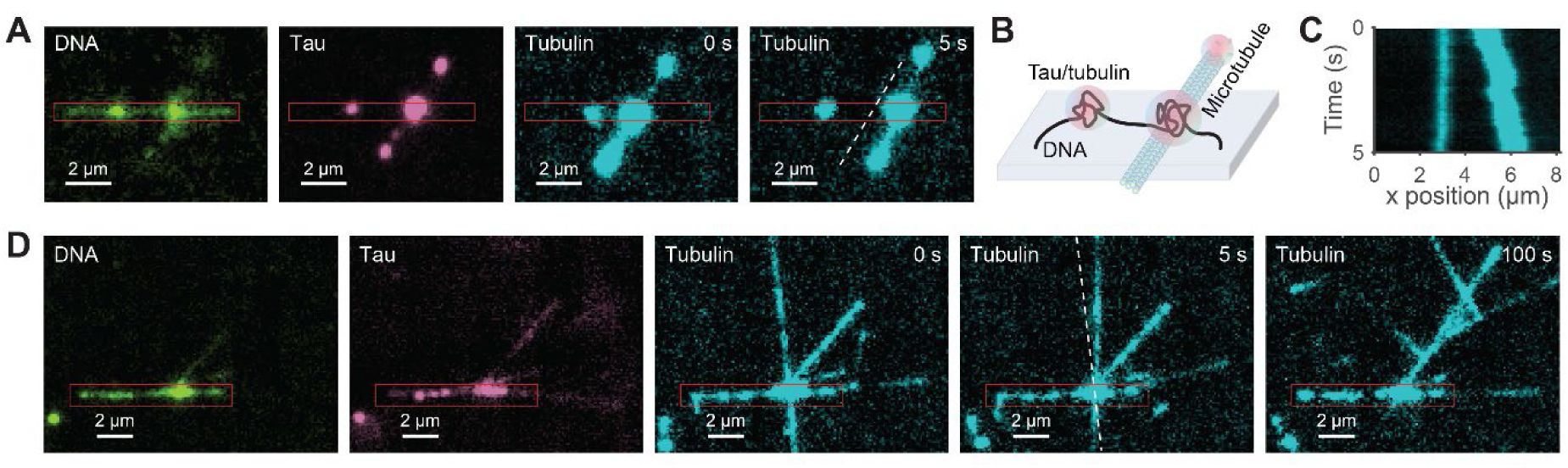
Microtubules interfacing with tau–DNA co-condensates (A) Snapshots showing the dynamic gliding of microtubules (*cyan*, labeled with HiLyte 488,) along DNA (*green*, stained with SYTOX Orange), driven by the mobility of tau condensates (*magenta*, labeled with Cy5) on DNA. Microtubules were formed by incubating with 5 μM tubulin. (B) Schematic illustration of the DNA–tau–microtubule tripartite assembly. (C) Representative kymograph displaying the mobility of the gliding microtubule on DNA shown in (A). (D) Time-lapse images capturing the pivoting motion of the microtubule, anchored by tau–DNA co-condensates. In (A) and (D), red boxes mark the location of the DNA strands, and white dashed lines show the position of the microtubules in the preceding image.

Intriguingly, microtubules attached to both singly and doubly tethered DNA strands exhibited dynamic behaviors, including diffusive movements along DNA due to mobile condensates (Fig. 4A–C, Movie S6) and pivoting around anchoring tau condensates (Fig. 4D, Movie S7). This was particularly evident in the occasional diffusive translation of entire microtubule filaments along double-tethered DNA substrates. With a slight increase in DNA surface density, more microtubules were captured, and tau clusters on different DNA strands cooperatively bridged and bundled nearby microtubules, forming a tripartite network of DNA, tau, and microtubules (Movie S8). These observations suggest that tau condensates may facilitate the repositioning and stabilization of microtubules relative to DNA, in processes such as mitotic spindle attachment to chromosomes.

To directly demonstrate the role of tau condensates in microtubule capture, we designed a microtubule capture assay in a more complex environment with higher concentrations of tau and DNA. In this assay, we densely packed the surface with DNA (Fig. 5A, Movie S9), allowing multiple strands to collaboratively induce tau condensation. We increased the tau concentration to 5 μM and added 5% PEG to simulate molecular crowding. The formation of tau–DNA co-condensates appeared as discrete bright nodes interconnected by surface-tethered DNA, which manifested as thin, dimmer lines forming a reticular network (Fig. 5B, Movie S9). The tau–DNA co-condensates appeared as distinct bright foci, while the surface-tethered DNA formed an intricate mesh connecting these nodes. This reticular structure suggests a complex and organized arrangement of tau–DNA interactions, highlighting tau’s ability to induce localized DNA condensates while maintaining a networked distribution across the nucleic acid surface, similar to the behavior of various DNA-compacting proteins. Under this condition, the overall structure became more stabilized, and the mobility of tau locations was significantly reduced (Movie S9).

**Fig. 5.**
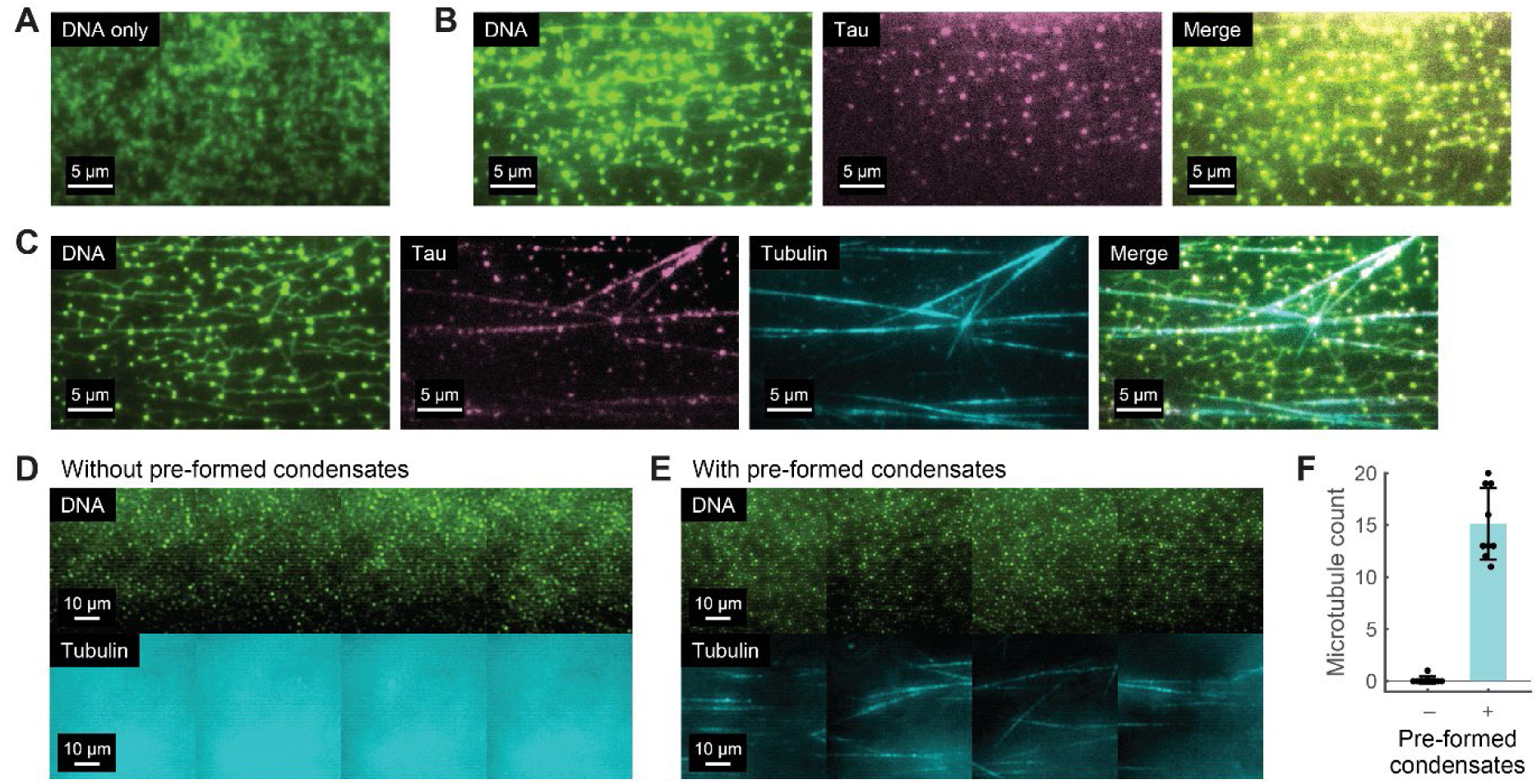
Microtubules captured by dense tau–DNA co-condensates (A–C) TIRF images of high-density surface-tethered λ-DNA (*green*, stained with SYTOX Orange) before tau addition (A), after adding 5 μM tau (*magenta*, Cy5-labeled) with 5% PEG (B), and after adding a mixture of tau and tubulin (*cyan*, HiLyte 488-labeled) with 5% PEG (C). Microtubules were pulled to the surface by tau–DNA co-condensates. (D, E) Large-area views of microtubule pulldown on DNA surfaces with and without pre-formed tau–DNA co-condensates. In (D), microtubules did not attach to the DNA surface without tau pre-incubation to form condensates; only background fluorescence from out-of-focus tubulins was visible. (F) Quantification of surface-bound microtubules with and without pre-formed tau–DNA co-condensates (*n* = 9 images; mean ± s.d.).

Under these conditions, tau–DNA clusters at distant sites were able to cooperatively recruit longer microtubules (Fig. 5C, Movie S9), effectively bridging the gap in length scale. Since the DNA molecules were arranged on a 2D plane, the captured microtubules conformed to the substrate surface. Most importantly, when tau condensates were not pre-formed before the introduction of tubulin, only a few surface-bound microtubules were observed (Fig. 5D), despite the presence of numerous microtubules away from the DNA surface (Movie S10), contributing to background fluorescence. By quantifying the number of microtubules attached to the DNA surface, we confirmed that the presence of pre-formed tau–DNA co-condensates is indeed critical for capturing microtubules from the solution. We speculate that this conformal attachment of microtubules to a densely packed DNA surface via tau–DNA co-condensates may represent a mechanism for docking nascent microtubules to complex DNA structures, such as mitotic chromosomes.

### Phosphomimetic tau mutants condense DNA but differ in microtubule-interfacing potential

Phosphorylation of tau at multiple sites is tightly regulated by various kinases and phosphatases, influencing its affinity for microtubules (*39*). To investigate how phosphorylation-like changes affect tau–DNA interactions and their role in recruiting microtubules, we generated two additional recombinant tau proteins with phosphomimetic mutations: T231D/S235D (a double mutant) and S262D. Given the critical association of these sites with microtubule binding and Alzheimer’s disease (*40–43*), these mutants are often used to model pathological modifications seen in tauopathies (*44–47*).

We first confirmed through single-molecule experiments that both T231D/S235D and S262D tau proteins successfully undergo co-condensation with DNA, similar to wild-type tau (Fig. 6A). Next, we examined whether these mutants could still support the assembly of the DNA–tau–microtubule network. When mixed with tubulin in the presence of GTP, the phosphomimetic mutants were able to nucleate microtubule growth, though to a much lesser extent than wild-type tau (Fig. 6B). Specifically, the microtubules formed in the presence of these mutants were shorter, less bundled (as indicated by a dimmer appearance), and depolymerized more rapidly compared to those formed with wild-type tau (Fig. S5). Nonetheless, in this buffer condition, microtubule growth was entirely absent without tau (Fig. S5), indicating that the tau mutants still retain some microtubule-binding activity. As a result, wild-type tau outperformed the phosphomimetic mutants in promoting microtubule growth, as reflected by the higher microtubule count (Fig. 6C). These findings are consistent with previous observations that phosphorylated tau has impaired microtubule binding, leading to a reduced stabilizing effect.

**Fig. 6.**
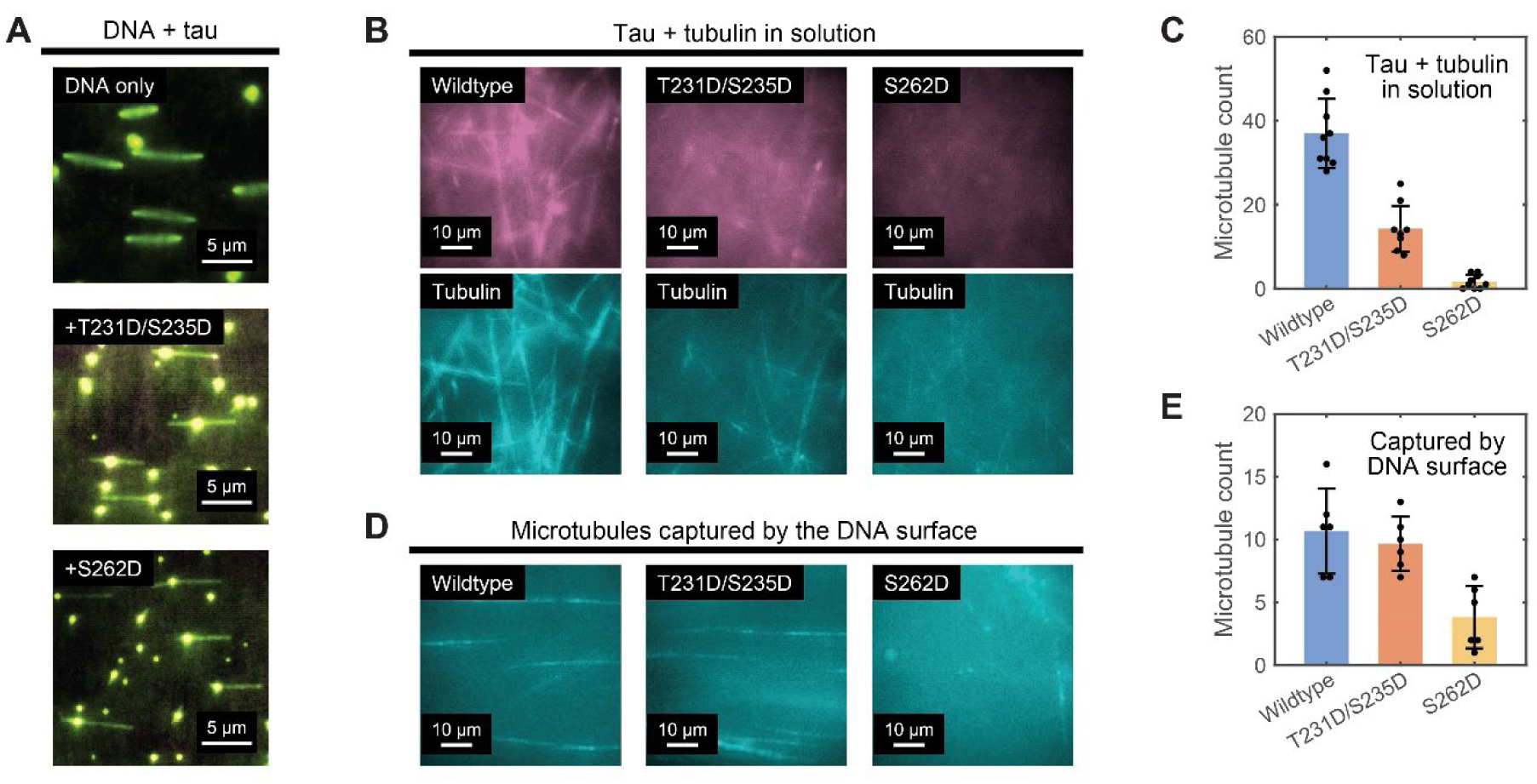
Phosphomimetic tau mutants in DNA condensation and microtubule interactions (A) TIRF images (time-averaged projection over 5 s) of DNA alone (*top*) and DNA incubated with the indicated tau mutants (T231D/S235D and S262D). (B) Microtubule formation in solution in the presence of the indicated tau species. (C) Quantification of microtubule counts in the tau–tubulin mixture (*n* = 9 images; mean ± s.d.), as shown in (B). (D) Large-area views of microtubule capture on the DNA surfaces with tau condensates formed by the indicated species. (E) Quantification of microtubule counts on the surface captured by DNA (*n* = 6 images; mean ± s.d.), as shown in (D).

To compare the capacity of different tau mutants to couple microtubules to DNA, we performed a microtubule capture assay using tau–DNA co-condensates, as previously described for wild-type tau (Fig. 5D, E). The images of microtubules on the DNA surface were largely indistinguishable between wild-type and the T231D/S235D mutant. However, in the case of the S262D mutant, almost no microtubules were observed; instead, clustered aggregates of tubulin formed (Fig. 6D). Quantification of the microtubule count revealed that, despite its impaired ability to promote microtubule formation in solution (Fig. 6B, C), the T231D/S235D tau mutant captured and interfaced with microtubules on the DNA surface at a level comparable to wild-type tau (Fig. 6D, E). This suggests that the T231D/S235D mutant retains a particularly strong capacity for mediating interactions between DNA and microtubules, even though it exhibits reduced microtubule nucleation and bundling overall. In contrast, the S262D mutant showed a marked decrease in microtubule capture on the DNA surface, presumably due to its impaired microtubule binding and stabilization (Fig. S5) (*41*). Overall, these results suggest that tau’s ability to interface between DNA and microtubules may be regulated by site-specific phosphorylation.

### Tau clusters localize on centromeres during prometaphase of mitosis

The tripartite assembly of DNA, tau, and microtubules observed in vitro led us to hypothesize their potential role in cell division, particularly in the early recognition of mitotic chromosomes by nascent mitotic spindles. To investigate tau distribution during cell division, we first used HEK-293 cells, which have low endogenous tau levels, and transiently expressed recombinant wild-type 2N4R tau tagged with mCherry. Tau expression caused slight transcriptomic changes, as shown by RNA-seq analysis (Fig. 7A). These changes included the upregulation of several stress-response genes, such as CHAC1, ATF3, HMOX1, and DDIT3, which are involved in the unfolded protein response, oxidative stress, and apoptosis. In addition, downregulation was observed in genes such as CENPE, which is involved in chromosome movement, and BRCA2, which plays a role in DNA damage repair and cell cycle regulation (Fig. 7B). These changes primarily affected genes related to cell division, microtubule organization, and DNA repair. However, the overall impact on cell proliferation in tau-expressing cells was minimal.

**Fig. 7.**
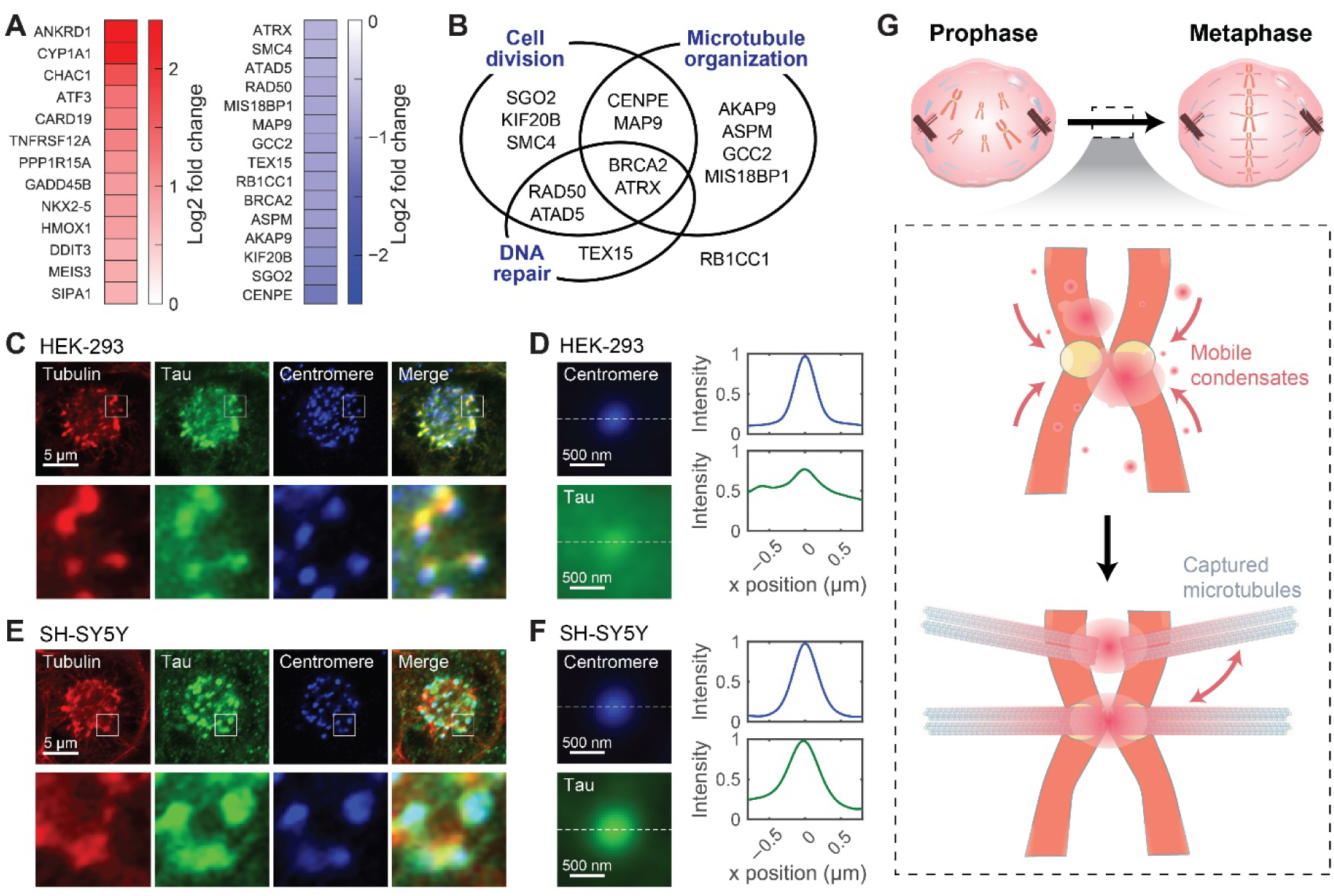
Localization of tau on centromeres during cell division (A) List of top up- (*left*) and down-regulated (*right*) genes in HEK-293 cells transiently expressing tau–mCherry (log2 fold change > 0.7 or < −0.7; *p* < 0.05). (B) Venn diagram for the downregulated genes shown in (A), involved in cell division, microtubule organization, and DNA repair pathways. (C) Immunofluorescence images of prometaphase tau-expressing HEK-293 cells showing the distribution of tubulin, tau, and centromeres. (D) Intensity profiles of centromere (*top blue*) and tau (*bottom green*) measured across aligned centromere regions. Centromere images were collected from multiple regions (*n* = 36 locations from 18 cells), aligned, and analyzed for the presence of tau signal. (E, F) Same as (C) and (D), except for the use of SH-SY5Y cells endogenously expressing tau (*n* = 22 locations from 11 cells for (F)). (G) A proposed model illustrating tau–DNA co-condensation during mitosis. In prometaphase, tau forms mobile condensates on chromosomes, aiding in the capture and dynamic rearrangement of nascent microtubules. As these condensates merge and migrate towards the centromere, stable chromosome-spindle connections are established via kinetochores.

Confocal imaging of HEK-293 cells immunostained for microtubules and centromeres, along with mCherry fluorescence from tau, was performed. We focused on cells in prometaphase, just before chromosome congression, by synchronizing them with a thymidine block. Images of cells in this phase displayed the characteristic rosette structure formed by mitotic chromosomes surrounding the nascent mitotic spindle (Fig. 7C). These images also revealed strong colocalization between tau and the newly forming mitotic spindles, indicating that tau remains closely associated with microtubules during prometaphase. Notably, tau formed distinct clusters on mitotic chromosomes, suggesting its involvement in chromosome-spindle attachment (Fig. 7C, magnified views in the bottom panels). To quantitatively assess the colocalization between tau and centromeres, we pooled images of 36 centromeres from 18 cells and averaged them after precise alignment (Fig. 7D). Intensity profiles of centromeres and tau fluorescence in these averaged images indeed confirmed colocalization, with tau intensity peaks overlapping with centromere signals.

To corroborate the physiological relevance of our findings, we next examined tau distribution in SH-SY5Y cells, a widely used neuroblastoma model that endogenously expresses tau. Through immunofluorescence imaging of synchronized SH-SY5Y cells using an antibody against tau, we captured prometaphase cells (Fig. 7E), yielding results similar to those observed in HEK-293 cells. Once again, we observed a high degree of colocalization between tau and microtubules. Additionally, tau formed clear foci on mitotic chromosomes, further supporting its role in chromosome-spindle attachment. Quantitative analysis of 22 centromeres from 11 cells also confirmed a clear overlap between tau and centromere markers (Fig. 7F). Collectively, these findings highlight the consistent involvement of tau in interfacing microtubules with centromeres during mitosis across different cell types, whether tau is expressed endogenously or exogenously.

## Discussion

Despite extensive research, many aspects of tau biology remain unclear, including its functions beyond microtubule stabilization and the precise mechanisms by which tau contributes to disease progression. Our results from single-molecule experiments conclusively demonstrate that tau induces local compaction of naked DNA, forming dynamic, mobile condensates (Fig. 2). This behavior resembles LLPS of tau in solution, which has been linked to the formation of insoluble tau aggregates, such as neurofibrillary tangles, in neurodegenerative diseases. However, the tau– DNA co-condensation we observe occurs below the critical saturation concentration (*c*sat) for tau alone (50–100 μM) or under crowding conditions (>1 μM) (*5*, *6*, *27*), suggesting that DNA strands enrich the local concentration of tau, reminiscent of tau interacting with RNA and RNA-binding proteins (*4*, *12–14*, *16*). DNA binding effectively reduces the dimensionality of tau clustering (from 3D to quasi-1D), potentiating condensation similarly to LLPS on 2D lipid membranes (*48*). Tau’s co-condensation with DNA is likely driven by a combination of electrostatic and hydrophobic interactions, again similar to its behavior in LLPS with RNA (*6*, *15*, *49*). Similar protein–DNA co-condensation behaviors are being discovered in other DNA-binding proteins (*28*, *31–34*), which can generate forces on DNA (*30*) as demonstrated in our magnetic tweezers experiments (Fig. 3). Capillary forces and dynamic instability are expected to regulate the formation and coalescence of these condensates (*50*, *51*), influencing their size, shape, and interactions with surrounding structures, including microtubules.

Given the contour length of dsDNA (∼0.34 nm/bp), the 50–70-nm increments observed during the stretching of tau–DNA co-condensates (Fig. 3F) correspond to 150–200 base pairs. This is intriguingly close to the persistence length of dsDNA (∼150 bp) and also the wrapped length in nucleosomes (147 bp), suggesting that tau condensates may associate with DNA segments of similar size. Each step likely represents the overcoming or release from a single tau cluster. The minor groove binding of tau to the DNA backbone (*20*) may sculpt this specific compaction, helping tau overcome the intrinsic stiffness of dsDNA and promoting the formation of more organized structures. Additionally, pre-formed tau clusters may guide the positioning of evolving condensates along certain DNA regions, contributing to the stepwise behavior. In contrast, the smoother changes during the condensing phase suggest a gradual accumulation of tau molecules with the DNA strands being steadily drawn into the ripening tau condensates. The rare stepwise events in this phase likely result from sudden compaction of DNA as it partitions into existing tau condensates. Further structural studies on tau–DNA complexes using cryo-electron microscopy or X-ray crystallography will be in order.

We also demonstrate that tau–DNA co-condensates can capture and organize microtubules, acting as dynamic hubs (Fig. 4). It is surprising that microtubules did not attach to DNA without pre-formed tau condensates (Fig. 5D–F), even when the microtubule solution contained the same concentrations of tau and PEG used during preincubation. This implies a distinct difference between tau condensates formed on DNA and those mixed with tubulin, which requires further investigation. The multivalent nature of tau likely enables its interaction with both microtubules and DNA, a process that may be fine-tuned by post-translational modifications such as phosphorylation and acetylation. These modifications, which adjust tau’s charge, may act as switches controlling its partitioning between microtubules and DNA (*52*). Notably, tau phosphorylation is an important mechanism for regulating microtubule stability and is controlled in a cell cycle-dependent manner (*53–56*). Given the differential effects of the site-specific phosphorylation we tested (Fig. 6), we suggest that this balance influences tau’s multifaceted interactions with mitotic spindles, chromosomes, and the dynamic interplay between the two.

Tau’s ability to capture and stabilize microtubules around DNA suggests its involvement in bridging nascent mitotic spindles to chromosomes during cell division, facilitating the random “search and capture” process (*57*). We speculate on a hypothetical function of tau in chromosome capture during mitosis (Fig. 7G): (i) in prometaphase, tau condenses on DNA substrates, including chromosomes; (ii) the dynamic condensates locally remodel DNA structures and facilitate the docking of microtubules undergoing rapid polymerization and depolymerization, predominantly through the registration of their lateral surfaces (*58*); and (iii) as the condensates mature toward centromeres, the premature lateral attachment of microtubules via tau is gradually taken over by the firm end-on attachments mediated by kinetochores for subsequent metaphase congression(*59*). The idea of tau’s involvement in chromosome-spindle linkage was first considered in the 1980s (*60*) but has not been thoroughly explored, with experimental evidence remaining limited. Although our model does not necessarily invoke the function of the nuclear fraction of tau (*26*), it is possible that nuclear tau, if it occupies centromeres before nuclear envelope breakdown, could guide the migration of tau droplets on other chromosomal sites toward the centromeres. The reported localization of nuclear tau at pericentromeric regions (*61*, *62*), along with recent findings of tau’s involvement in heterochromatin and HP1α clustering (*35*, *63*, *64*), supports this idea. Note that this model is also compatible with other mechanisms that facilitate chromosome capture and biorientation through alternative pathways (*38*, *65–67*).

Tau’s association with centromeres during prometaphase, as evidenced by our imaging data from both HEK-293 cells (with transiently expressed tau) and SH-SY5Y cells (showing endogenous tau distribution) (Fig. 7C–F), supports the potential link between tau–DNA co-condensation and cell division. However, more evidence is required for a stringent test of this hypothesis, including determining the direct binding between tau and DNA on centromeres, and assessing the effects of tau depletion or mutations that specifically interfere with tau–DNA interactions. The transcriptomic shift observed in HEK-293 cells upon tau expression (Fig. 7A, B) likely reflects a mild cellular response to excess tau (*68–70*), consistent with its expected influence on microtubule dynamics, but may also point to additional roles in DNA protection (*24*, *35*, *71*, *72*) or broader signaling pathways (*73*). However, these shifts did not significantly disrupt cell division, allowing us to examine tau during mitosis without inducing excessive cellular stress.

As many cell types do not constitutively express tau, it is possible that the proposed mitotic action of tau–DNA co-condensation comes into play only under conditions of elevated tau levels in proliferating cells. In this case, when the balance of tau partitioning between DNA and microtubules is disrupted, tau may either fail to bind to chromosomes or bind excessively, akin to its wetting of microtubules. This disruption can interfere with chromosome segregation, potentially leading to stray chromosomes and aneuploidy. It is tempting to postulate that hyperphosphorylation of tau, long associated with Alzheimer’s disease and other tauopathies (*39*, *74*, *75*), may also contribute to chromosome missegregation through disrupted tau–DNA co-condensates. Given the link between aberrant cell cycle activation in postmitotic neurons and neurodegeneration (*76–78*), as well as the reduction of adult hippocampal neurogenesis in cognitive decline (*79*, *80*), it is plausible that defective tau function during cell division could have broader implications for neurodevelopment and neurodegeneration. We further contemplate the potential connection between our findings and tau expression in cancer (*81–84*), where disrupted mitotic signaling may provoke dysregulated tau activity during cell division, leading to chromosome instability and ultimately aneuploidy, a hallmark of cancer. Future studies on tau diversity, including developmental isoforms, pathogenic mutations, and phosphorylation status— which can alter its LLPS behavior and/or microtubule dynamics (*5*, *85–88*)—will explore how these changes impact tau–DNA co-condensation and the subsequent DNA–tau–microtubule assembly.

## Conclusion

We report that single-molecule experiments revealed strong, dynamic co-condensation between tau and naked DNA. These co-condensates can interface microtubules with DNA in vitro, and phosphomimetic tau mutants exhibited varying abilities in this function. Cell imaging indicated tau localization at centromeres during early mitosis, suggesting a role in chromosome capture and congression. Our findings introduce a new dimension to tau’s functionality, suggesting a potential connection to chromosomal dynamics during cell division. These insights may provide a foundation for understanding tau’s involvement in physiological and pathological processes beyond the nervous system. Further studies are needed to determine whether the proposed model holds true across various contexts and to explore how tau phosphorylation regulates its mitotic functions and contributes to disease when these processes are disrupted.

## Materials and Methods

### Protein purification and labeling

The 2N4R isoform of 6×His-tagged human tau protein was cloned into the pRK172 vector and expressed in LEMO21 cells (NEB #C2528J). Cells were cultured in TB media at 37 °C until the OD600 reached 0.6, followed by treatment with 10 mM betaine and 500 mM NaCl. After a 30-min incubation at 30 °C, 0.4 mM IPTG was added, and the culture continued for an additional 3–4 h. The cells were homogenized and lysed in lysis buffer (20 mM MES (pH 6.0), 0.2 mM MgCl2, 1 mM EGTA, 1.5 mM PMSF, 1× protease inhibitor cocktail, 5 mM DTT). The lysate was sonicated at 30% amplitude in 30-s on-off cycles for 3 min, followed by the immediate addition of 500 mM NaCl. The lysate was boiled for 15 min and centrifuged at 15,000 *g* for 1 h. Supernatants were purified using Ni-NTA resin (Qiagen). The bound resin was washed with wash buffer (50 mM phosphate buffer (pH 7.2), 200 mM NaCl, 20 mM imidazole) and eluted with elution buffer (50 mM phosphate buffer (pH 7.2), 200 mM NaCl, 500 mM imidazole). TEV protease (Sigma) was then added to remove the His-tag, and the solution was incubated overnight at 4 °C. To further purify the protein, ion exchange chromatography (Bio-Rad #1580030) was performed to remove dimers and other undesired proteins. The protein elutes were concentrated using a protein concentrator (Sartorius #VS15T01). For protein labeling, the purified protein was mixed with a 10-fold molar excess of Cy5-maleimide (Cytiva #25800674) and incubated for 2 h at room temperature (RT) in reaction buffer (50 mM phosphate buffer, 0.05 mM TCEP). Free dye was removed by gel filtration (Bio-Rad #1500738). The Cy5-labeled tau proteins were then concentrated using a protein concentrator (Sartorius #VS15T01).

### Fluorescence imaging of tau–DNA co-condensation

To prepare DNA constructs for surface attachment, λ-DNA (Thermo #SD0021) was mixed with a 100-fold molar excess of 5′- and 3′- biotinylated oligos and incubated in 1× NEB ligase buffer at 65 °C for 5 min. The mixture was then slowly cooled over 3 h. T4 DNA ligase (NEB #M0202L) and 1 mM ATP were added, followed by overnight ligation. The sample was purified using a 100 kDa cutoff buffer exchange filter (Merck, #UFC510024) to remove free primers and exchange the buffer for distilled water. For TIRF microscopy, a flow chamber was constructed using previously described methods (*89*, *90*), with meticulous passivation of the glass coverslip surface via PEGylation. After the first round of PEGylation using a 99:1 mixture of 5-kDa mPEG and biotin-PEG (Laysan Bio), the coverslips were subjected to a second round of overnight PEGylation with a solution of 2-kDa mPEG (Laysan Bio) without biotin-PEG to ensure optimal passivation. All experiments were performed on a home-built TIRF microscope setup, with sample solutions injected using a syringe pump. For capturing DNA samples, 8 nM NeutrAvidin (Thermo #31000) was introduced into the flow cell, incubated for 5 min, and then washed with PBS. The flow cell surface was further passivated by infusing 5% BSA (Sigma #A0100-010) for 10 min, followed by another PBS wash. Biotinylated λ-DNA was diluted to 10–20 pM, injected, and immobilized on the surface for 7 min. For imaging, the buffer was exchanged with 200 μL of imaging buffer (50 mM HEPES (pH 7.1), 100 mM NaCl, 2 mM Trolox, 59.5 nM protocatechuate 3,4-dioxygenase (PCD), and 2.5 mM protocatechuic acid (PCA)) containing 20 nM SYTOX Orange (Thermo #S11368) and incubated for 10 min. To examine tau–DNA co-condensation, tau protein was diluted in the imaging buffer to concentrations ranging from 50 nM to 5 μM and then injected into the flow cell. Samples were illuminated with 488 nm, 532 nm, and 633 nm lasers and imaged using a 60× oil-immersion lens (Olympus) and a high-sensitivity sCMOS camera (Photometrics Prime BSI Express), typically operated with a 100-ms exposure time using custom LabVIEW software. When necessary, fluorescence in two channels was separated by a dual-view image splitter (Cairn OptoSplit II) and recorded simultaneously. For the DNA skybridge experiments, a flow chamber was constructed using microfabricated quartz and a glass coverslip, as detailed in the Supplementary Materials and a previous publication (*36*). For sample assembly, 30 pM of biotinylated λ-DNA was injected into the flow chamber at a rate of 70 μL/min using a syringe pump (Harvard Apparatus) and tethered to both ridges of the quartz via NeutrAvidin. All imaging was conducted on a prism-type TIRF microscope (Olympus IX-71) equipped with a 60× water-immersion objective (NA = 1.2). SYTOX Orange-labeled DNA was imaged with 561-nm illumination, while Cy5-labeled tau was imaged with a 638 nm laser. Fluorescence images (100 ms per frame) were captured by an EMCCD camera (Andor iXon3 897) controlled by Micro-Manager imaging software.

### Magnetic tweezers experiments

The preparation of the 10-kbp dsDNA construct used in this work is detailed in our previous work (*37*). Briefly, three DNA fragments were generated by PCR from λ-DNA: one 10 kbp fragment and two 500 bp fragments (the list of primers is provided in the Supplementary Methods). During PCR, the 500 bp fragments were labeled with multiple biotin or digoxigenin molecules by incorporating biotin- or digoxigenin-labeled dUTP (Sigma). All three PCR products were digested with BsaI for 4 h at 37 °C, and after cleanup, were ligated overnight using T4 DNA ligase at RT. The ligated product was then purified by agarose gel electrophoresis. Details of the flow cell construction, sample assembly, and the home-built magnetic tweezers setup are available in our previous work (*91*, *92*). For measurements, the DNA samples were injected into the flow cell so that the biotinylated ends could attach to PEGylated glass surfaces via biotin-avidin interactions, while the digoxigenin-labeled ends were coupled to anti-digoxigenin-coated magnetic beads (Invitrogen M-270), which were injected afterward. To apply force, an antiparallel pair of cubic neodymium magnets (N52, 3/8′′) was placed above the flow cell, with a 1-mm gap between the magnets. The height and rotation of the magnets were controlled using a translation stage (PI, L-40620DD10) and a stepper motor. Magnetic beads were illuminated with a 680 nm super-luminescent diode and imaged using a 100× objective lens (Olympus, NA 1.45) with a high-speed CMOS camera (Mikrotron EoSens 3CXP). Before measurements, the integrity of the dsDNA constructs (i.e., the absence of nicks) was checked by rotating the samples and observing supercoiling behavior (Fig. S4). When necessary, the force-extension curves were corrected for the off-axis tilting of magnetic beads (*93*). For fluorescence verification, DNA-bound tau protein labeled with Cy5 was imaged simultaneously with magnetic tweezing using a 532 nm laser aligned for TIRF illumination and an sCMOS camera (Photometrics Prime BSI Express) (*37*). All measurements were conducted in a buffer containing 50 mM HEPES (pH 7.1), 100 mM NaCl, 59.5 nM PCD, 2.5 mM PCA, and 0.1% BSA.

### Microtubule capture assay

Tau protein was diluted to 5 μM in imaging buffer (same as above) supplemented with 5% 10 kDa PEG, and then slowly injected into a sample with biotinylated λ- DNA to form condensates. After co-condensate formation, HiLyte 488-labeled tubulin (Cytoskeleton, # TL488M-A) was mixed with unlabeled tubulin (Cytoskeleton, #T240-B), making up 4% of the total tubulin concentration. The mixture was prepared in imaging buffer supplemented with 1 mM GTP and 5% 10-kDa PEG to achieve a final tubulin concentration of 5– 10 μM, while maintaining the tau concentration from the co-condensation experiments. After microtubule polymerization, the solution was slowly injected (7 μL/min) into the chamber, with or without tau–DNA co-condensates.

### Immunofluorescence imaging and analysis

The list of antibodies used for immunofluorescence imaging is provided in the Supplementary Materials. HEK-293 cells expressing 2N4R wild-type tau tagged with mCherry were prepared via reverse transfection using polyethyleneimine (Sigma). A total of 2.5 × 10^5^ HEK-293 cells were seeded onto poly-L-lysine (PLL)-coated 35 mm glass-bottom confocal dishes (SPL) with 2 μg/mL tau–mCherry plasmids and 4 μg/mL polyethyleneimine (PEI) in DMEM (Sigma) supplemented with 10% FBS (Welgene), and incubated for 36 h. For SH-SY5Y cells, 1 × 10^6^ cells were seeded onto the same dishes in RPMI-1640 (Sigma) supplemented with 10% FBS and were incubated for 24 h. To synchronize the cells, they were treated with 2 mM thymidine for 16 h to induce the first synchronization block, then released into their respective media (with 10% FBS) for 8 h. This was followed by a second 16-h thymidine block. After the second block, the cells were washed with PBS and released into their specific media with 10% FBS. Cells in the M phase were collected 9 h after release. The cells were then fixed with ice-cold methanol for 10 min, permeabilized with 0.25% Triton X-100 in PBS for 15 min, and blocked with 5% BSA in PBS with 0.05% Tween 20 (PBST) for 1 h at RT. Primary antibodies were diluted in PBST and incubated for 1 h at RT or overnight at 4 °C. After washing in PBST, cells were incubated with secondary antibodies diluted in PBST for 1 h at RT. After antibody staining, the cells were washed in PBST and incubated with 1 mg/mL Hoechst 33258 (Sigma) diluted 1:1000 in PBST for 10 min at RT. Following a final wash in PBS, coverslips were mounted on confocal dishes using mounting solution (Dako). Finally, the cells were imaged using a Zeiss LSM 900 confocal microscope equipped with Airyscan 2.

### RNA sequencing and analysis

RNA was extracted from tau–mCherry-expressing HEK-293 cells and untreated HEK-293 cells for control. The extracted RNA was purified and sequenced using the NovaSeqX platform at Macrogen (Seoul, Korea). Paired-end raw sequencing reads were pre-processed with FastQC and trimmed using Trimmomatic-0.39. The cDNA reads were aligned with HISAT2.1, and differential expression was defined and quantified using the DESeq program with criteria of log2 fold change >0.7 or <−0.7, along with *p*-values <0.05. To identify biological pathways associated with the differentially expressed genes, ShinyGO 0.80 was used for analysis with the Gene Ontology (GO) database. For the heatmap, log2 fold changes of genes corresponding to specific pathways were plotted, and a Venn diagram was generated for pathways involving downregulated genes.

## Author Information

## Acknowledgments

We thank Yoori Kim, Jeong-Mo Choi, and Cherlhyun Jeong for helpful discussions; Euddeum E. Jeong, Gyu Ri Kim, Jiwon Jang, Sejoo Jeong, and Jong-Chan Lee for the help with cell imaging; Minkwon Cha, Sang-Hyeok Jeong, and Haeun Yoo for the help with experiments; and the members of the Shon laboratory for discussions and assistance. This work was supported by the National Research Foundation of Korea (NRF) grant funded by the Korea government (MSIT) (NRF-2022R1C1C1012176 and RS-2023–00218927).

## Author Contributions

M.J.S. conceived the project. C.P., J.J., Y.H., and M.J.S. designed the project. C.P., J.J., Y.H., C.L.-E., and S.-H.R. prepared samples. C.P. performed TIRF imaging and microtubule experiments. J.J. and S.-H.R. performed magnetic tweezers experiments. J.J. and C.L.-E. performed confocal imaging and bulk LLPS experiments. K.Y. conducted skybridge experiments and J.S. analyzed data. A.J. and S.H. carried out RNA sequencing and analyzed data. M.J.S., D.S.H., and J.-B.L. secured funding and supervised the study. C.P., J.J., J.S., and S.H. prepared the figures and wrote the first draft. M.J.S., C.P., and J.J. wrote and edited the manuscript, with input from all authors.

## Competing Interests

None declared.

## Notes

### Competing Interest Statement

The authors have declared no competing interest.

