## Supplementary Materials for "Tau condensation on DNA and localization on centromeres: A potential link to cell division"

Park *et al.*

### The PDF file includes:

Supplementary Methods

Supplementary Figures S1–S5

Supplementary Video Legends

### Supplementary Methods

#### Electrophoretic mobility shift assay (EMSA)

DNA constructs were prepared either by PCR or annealing, using the primers and oligos listed below. The 5‑kbp and 500‑bp constructs were produced by PCR with nTaq polymerase from λ-DNA. A 24‑bp duplex was produced by annealing two complementary oligos, heating them to 95 °C, and slowly cooling to room temperature. To examine the interaction between tau and DNA, each construct was mixed with 50 nM – 5 μM 2N4R WT tau in a buffer containing 50 mM HEPES (pH 7.1) and either 25 or 100 mM NaCl, followed by a 1‑h incubation at room temperature. The mixtures were then run on a 1% agarose gel in 1× Tris-acetate-EDTA for 30 min at 100 V and stained with SYBR Safe DNA gel stain (Thermo).

#### Bulk LLPS and co-condensation of tau and DNA

Tau–DNA co-condensation via LLPS in bulk was examined in the presence of 10% PEG (#92897, Sigma, PEG 10,000) as a molecular crowding agent. A mixture of 0.1–10 ng/μL DNA constructs and 1–10 μM tau proteins, including 1% Cy5-labeled tau, was prepared in a buffer containing 50 mM HEPES (pH 7.1), 100 mM NaCl, 5.9 nM PCD, 2.5 mM PCA, and 20 nM SYTOX Orange. After a 1‑h incubation at room temperature, the samples were injected onto a PEGylated glass surface. The resulting condensates were imaged using a TIRF microscope equipped with a 60× oil-immersion objective.

#### Preparation of patterned quartz slides for DNA skybridge

Piranha-cleaned quartz slides were patterned as described in a previous study (Kim et al., 2019; doi: 10.1093/nar/gkz625). Briefly, ridge patterns were created using photolithography with TDMR-AR87 photoresist followed by HF etching. The final slides had a ridge interval of 13.1 μm, a barrier height of 4 μm, and a barrier width of 0.5 μm. For passivation, the quartz slides were rigorously cleaned with 10% Alconox, acetone, 1 M potassium hydroxide, piranha solution, and methanol, followed by brief flame exposure. The slides were then passivated with PEG, similar to the method described in the main text. NeutrAvidin was applied to the ridges by incubating a piece of PDMS (made from SYLGARD® 184, Dow Corning) with a NeutrAvidin solution (1 mg/mL in PBS) for 30 min before applying it to the patterned slides. The stamped slides and a glass coverslip were assembled with double-sided tape to construct a flow chamber for imaging.

#### List of PCR primers and oligos

- Oligos for the biotinylation of λ-DNA (for fluorescence imaging):
  - 5′-(Phosphorylation) AGG TCG CCG CCC TT (Biotin)-3′
  - 5′-(Phosphorylation) GGG CGG CGA CCT TT (Biotin)-3′
- Oligos for the 24-bp construct (for EMSA and bulk LLPS):
  - 5′-TCG ACA CGG AAA TGT TGA ATA CTA-3′
  - 5′-TAG TAT TCA ACA TTT CCG TGT CGA-3′
- Primers for PCR of the 500-bp construct (for EMSA and bulk LLPS):
  - 5′-ACG CGA AAA AAT GCC TGG-3′
  - 5′-TCG CCA CCA TCA TTT CCA-3′
- Primers for PCR of the 5-kbp construct (for EMSA and bulk LLPS):
  - 5′-TTA GAG AGT ATG GGT ATA TGA CAT CG-3′
  - 5′-TCG CCA CCA TCA TTT CCA-3′
- Primers for PCR of the 10-kbp construct (for magnetic tweezers experiments):
  - 5′-TTT TTT GGT CTC TAG ACT GAA CTT AAC GGG GCA TCG TA-3′
  - 5′-TTT TTT GGT CTC TCA CAC AAA AAT AAA GGG AAA GAT AAG CGC T -3′
- Primers for PCR of the 500-bp construct (for magnetic tweezers experiments):
  - 5′-TTT TTT GGT CTC TTG TGG TTC TGG CGT CGT TCT CG-3′
  - 5′-CGG AAC GCG ACA TCA CTC-3′

The underlines represent extensions for BsaI cuts.

#### List of antibodies for immunofluorescence imaging

- For HEK-293 cells expressing tau–mCherry:
  - Human anti-CREST (#15-234, Antibodies Incorporated, 1:200 dilution)
  - Mouse anti-alpha tubulin (clone DM1A, #62204, Invitrogen, 1:200 dilution)
  - Donkey anti-human Alexa Fluor 488 (#709-545-149, Jackson ImmunoResearch)
  - Goat anti-mouse Alexa Fluor 647 (#A-21235, Invitrogen)
- For SH-SY5Y cells with endogenous tau:
  - Human anti-CREST (#15-234, Antibodies Incorporated, 1:200 dilution)
  - Rabbit anti-alpha tubulin (#AB18251, Abcam, 1:200 dilution)
  - Mouse anti-Tau-1 (clone PC1C6, #MAB3420, Sigma, 1:200 dilution)
  - Donkey anti-human Alexa Fluor 488 (#709-545-149, Jackson ImmunoResearch)
  - Goat anti-rabbit Alexa Fluor 647 (#A-21244, Invitrogen)
  - Goat anti-mouse Alexa Fluor 568 (#A-11004, Invitrogen)

### Supplementary Figures

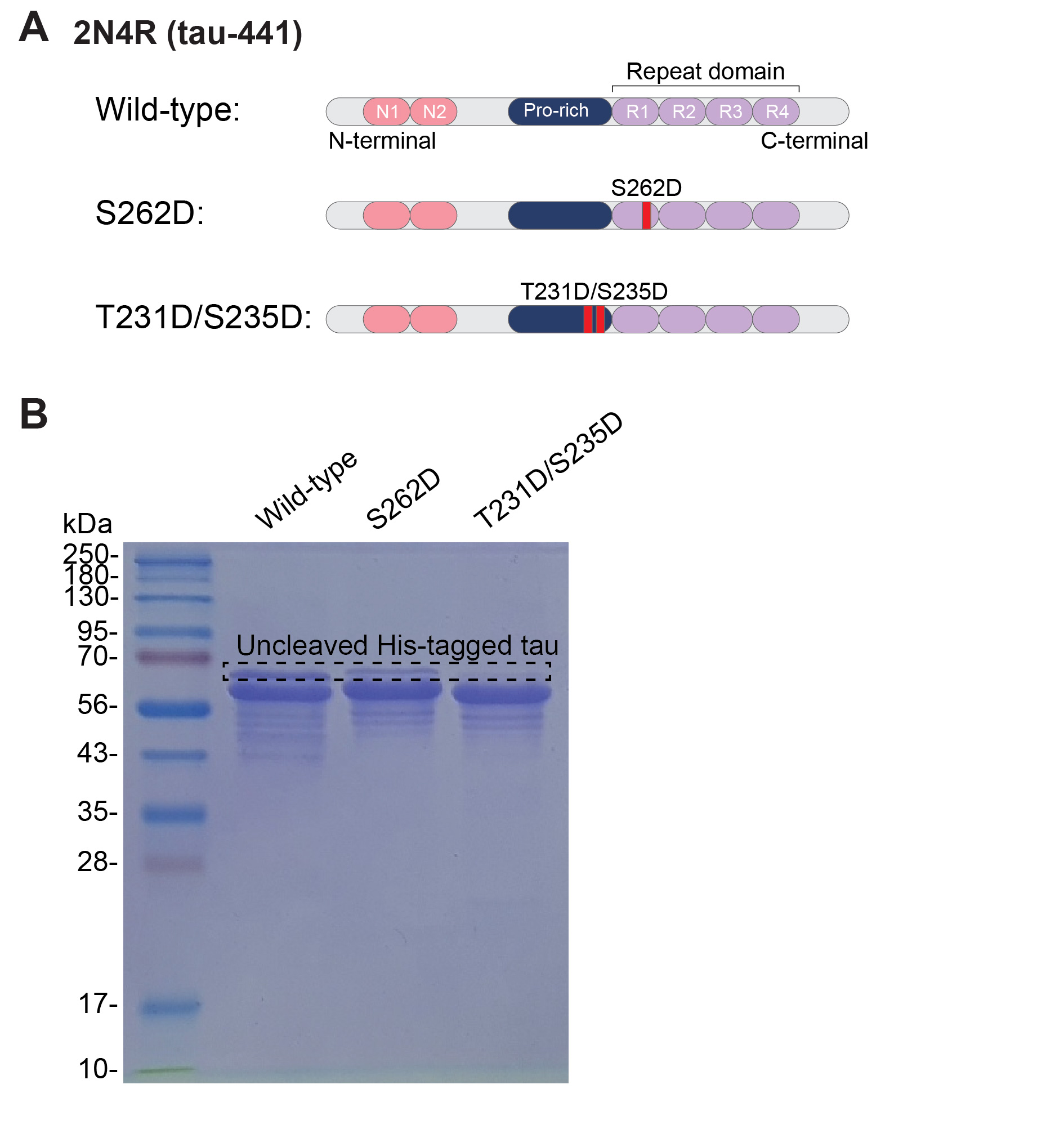

Figure S1. Recombinant tau proteins used in this study

(A) Domain structure of human 2N4R tau (tau-441), highlighting the sites of phosphomimetic mutations (S262D and T231D/S235D). (B) SDS-PAGE gel visualization of the purified recombinant proteins.

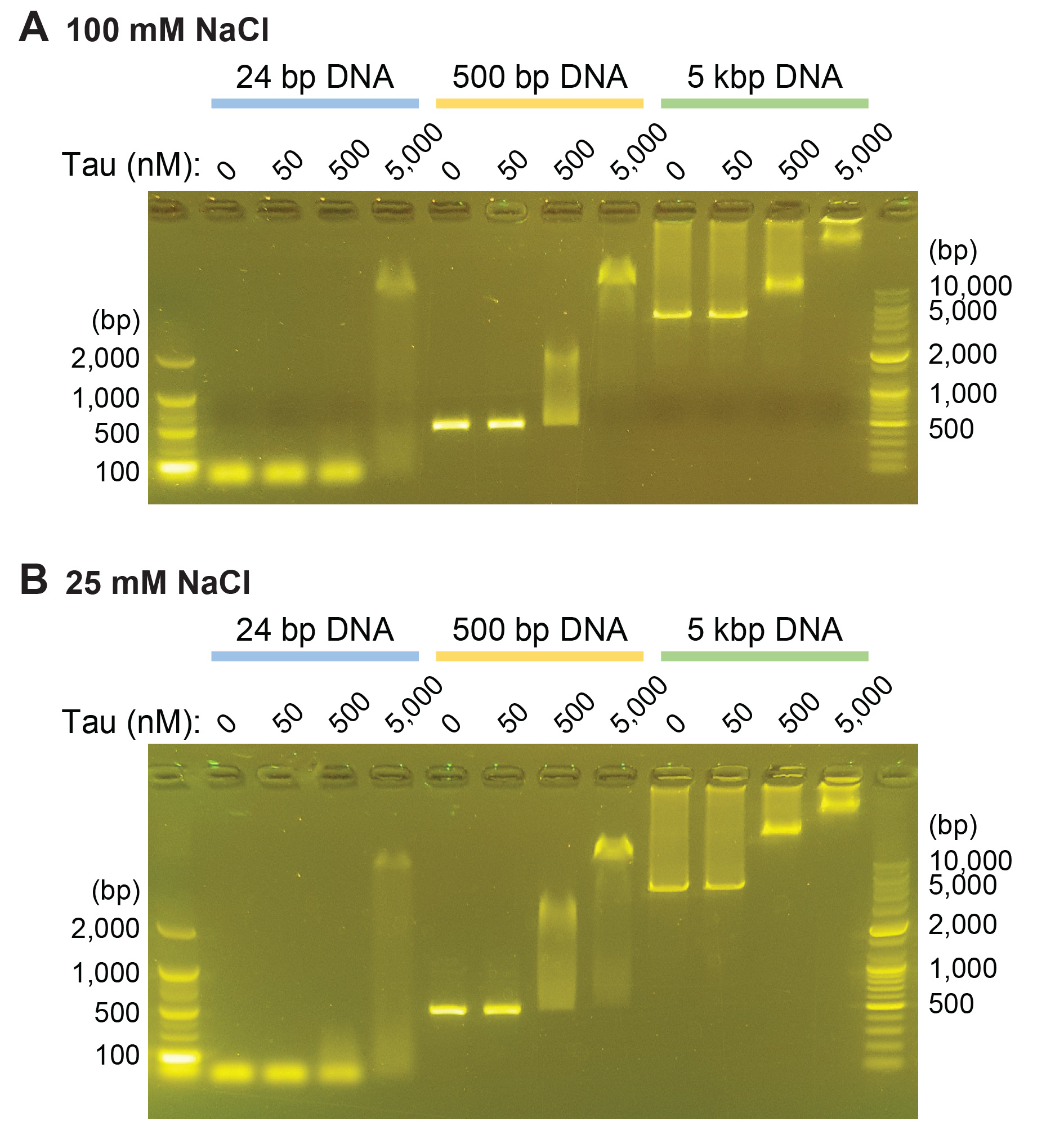

Figure S2. Electrophoretic mobility shift assay for tau–DNA interaction

Tau and DNA samples prepared under the indicated conditions were incubated for 1 h and run on a 1% agarose gel (see Supplementary Methods). Interactions were examined under (A) 100 mM NaCl and (B) 25 mM NaCl.

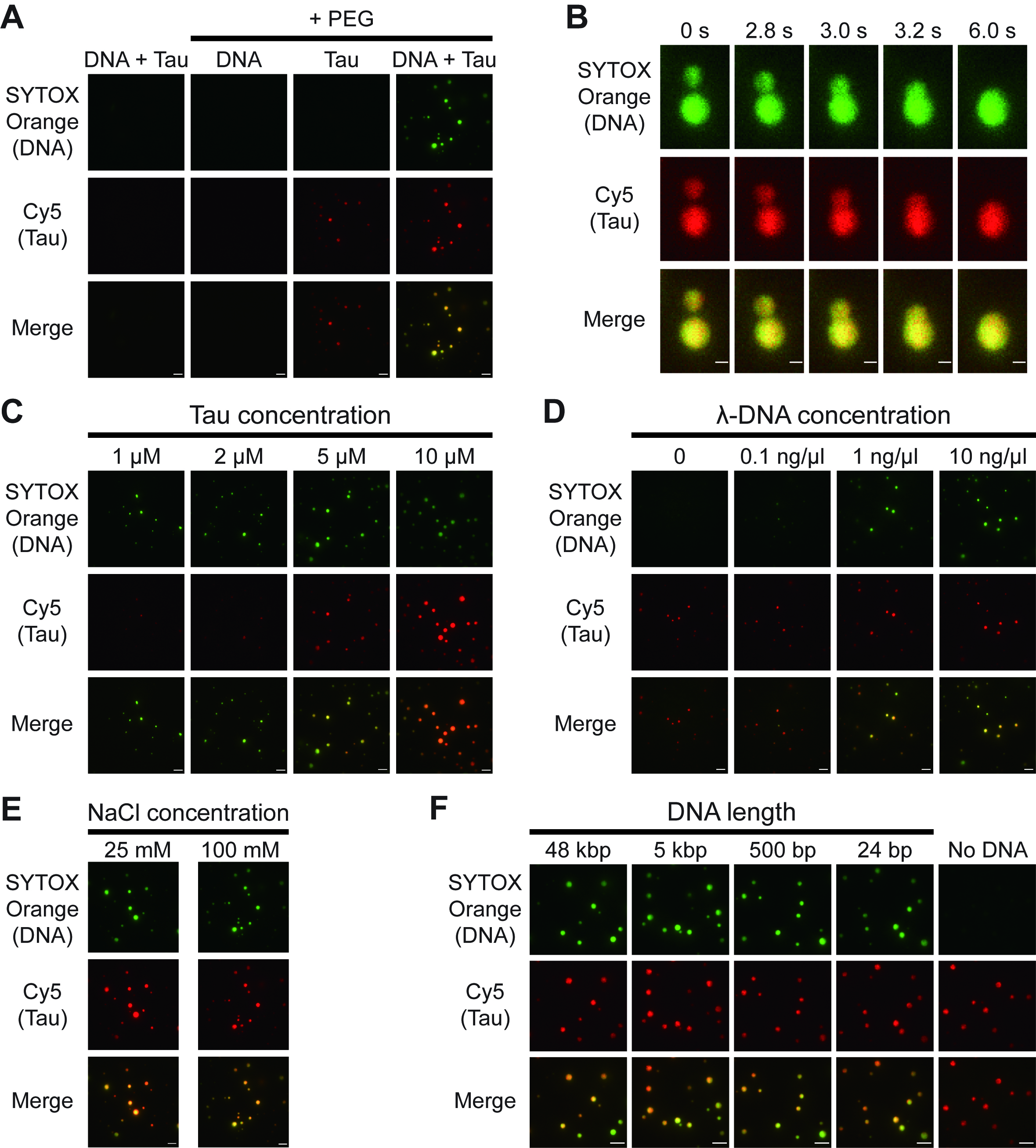

Figure S3. Co-condensation of tau and DNA in bulk via LLPS

(A) Co-condensation of tau (5 μM) and λ-DNA (10 ng/μL) observed with 10% PEG. (B) Merging of two co-condensates, characteristic of liquid droplets. (C-E) Effects of tau, DNA, and salt concentrations, with λ-DNA and tau kept at 10 ng/μL and 5 μM respectively. (F) Effect of DNA length, with 1 ng/μL DNA and 5 μM tau. Scale: 2 μm in (B), 5 μm for the rest.

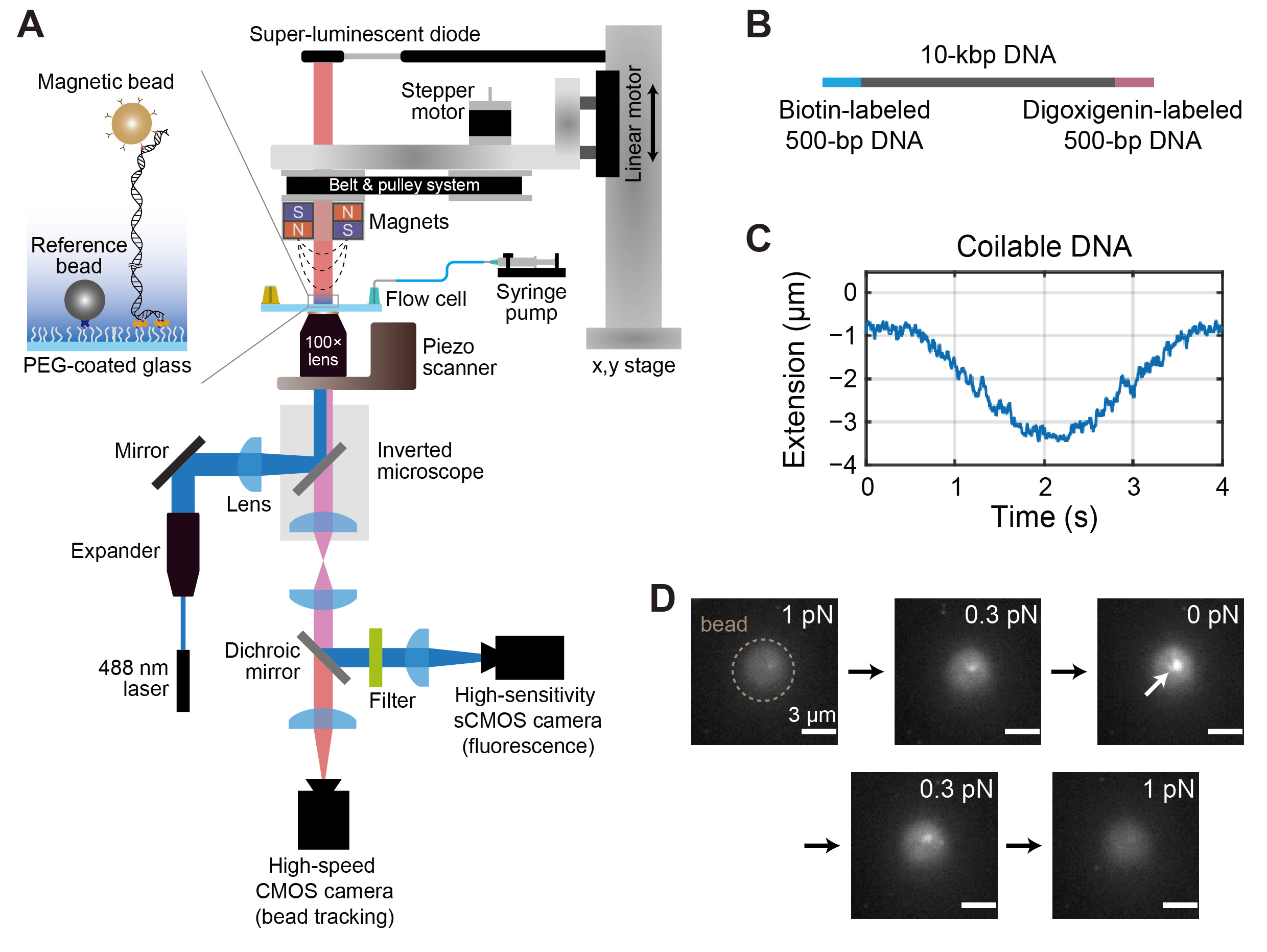

Figure S4. Verification of single-molecule constructs in magnetic tweezers experiments

(A) Schematic of the magnetic tweezers setup equipped with a TIRF imaging module. The inset is zoomed in to show the magnetic bead attached to DNA and the reference bead anchored to the surface. (B) Structure of the 10-kbp dsDNA construct, with two 500-bp fragments used for attachment to the surface and bead. (C) Verification of the intact dsDNA construct without nicks, demonstrated by its supercoiling behavior upon torque application via magnetic bead rotation (at 0.3 pN). (D) Fluorescence imaging of the bead–DNA construct during a force ramp (0.1 pN/s), using TIRF illumination in the presence of 5 μM tau, of which 50 nM was labeled with Cy5. The dashed circle marks the location of the magnetic bead with autofluorescence background, and the arrow points to the DNA tether with accumulating tau fluorescence. See also Movie S5.

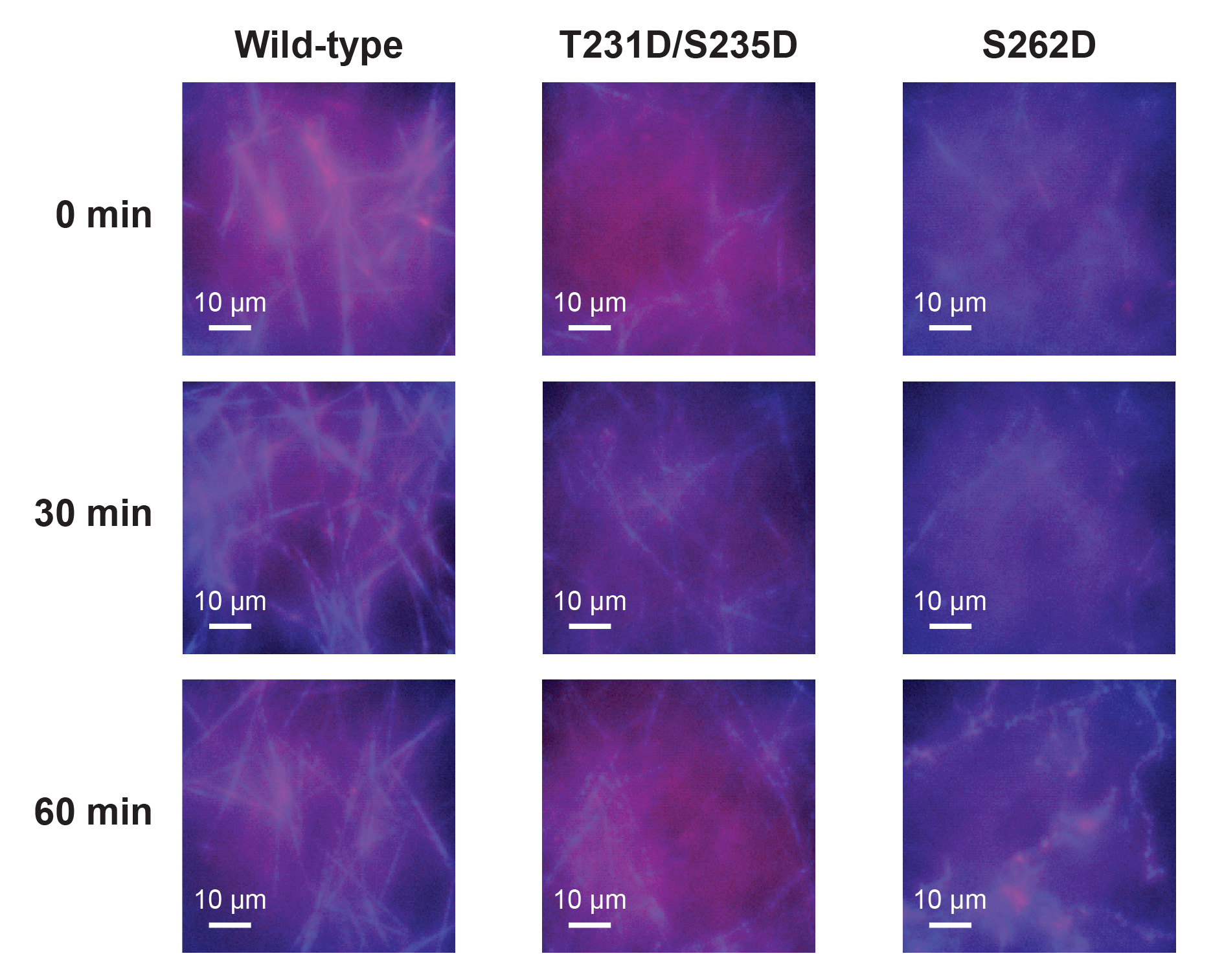

Figure S5. Stability of microtubules formed with different tau species

Microtubule stability over time in the presence of 5 μM tau of the indicated species, GTP, and 5% PEG. Images were captured at 0, 30, and 60 min to assess the structural integrity of the microtubules. Microtubules formed with wild-type (2N4R) tau remained very stable for at least 1 h, while T231D/S235D produced fewer microtubules, and those with S262D gradually decayed and aggregated. Blue: HiLyte 488-labeled tubulin; Red: Cy5-labeled tau.

### Supplementary Video Legends

Movie S1. Condensation of single-tethered λ-DNA by tau

A video of single-tethered λ-DNA under flow stretch, with condensation monitored during the introduction of 500 nM tau solution. Green: SYTOX Orange-labeled DNA; Magenta: Cy5-labeled tau. Exposure time: 0.1 s.

Movie S2. Condensation of double-tethered λ-DNA by tau

A video of double-tethered λ-DNA fluctuations in the presence of varying concentrations of tau and NaCl. Different molecules were imaged separately. Green: SYTOX Orange-labeled DNA; Magenta: Cy5-labeled tau. Exposure time: 0.1 s.

Movie S3. Mobility of tau-DNA co-condensates in surface-tethering experiments

A video of tau–DNA co-condensates moving along the double-tethered λ-DNA strand on the PEG-coated glass surface. Green: SYTOX Orange-labeled DNA; Magenta: Cy5-labeled tau. Exposure time: 0.1 s.

Movie S4. Mobility of tau-DNA co-condensates in skybridge experiments

A video of tau–DNA co-condensates moving along the λ-DNA strands prepared on the DNA skybridge platform. λ-DNA was stained with SYTOX Orange and imaged with 561‑nm illumination, while tau was labeled with Cy5 and imaged with 638‑nm illumination. The white box marks the region expanded on the right for a single condensate. Exposure time: 0.1 s, playback sped up 10×.

Movie S5. Verification of tau fluorescence in magnetic tweezers experiments

A video of fluorescence in the red channel for monitoring the accumulation of Cy5-tau during force ramp application in magnetic tweezers. Snapshots from this video are presented in Fig. S4D in the main text. Recorded at 0.1 s/frame and averaged over 1 s, with playback sped up 10×.

Movie S6. Translational movement of microtubules on tau-DNA co-condensates

A video of a microtubule moving along a DNA strand via tau condensates. Snapshots from this video are presented in Fig. 4A in the main text. Green: SYTOX Orange-labeled DNA; Magenta: Cy5-labeled tau; Cyan: HiLyte 488-labeled tubulin. The three channels were not recorded simultaneously. Exposure time: 0.1 s.

Movie S7. Rotational movement of microtubules on tau-DNA co-condensates

A video of microtubules attached to DNA via tau condensates (not shown), pivoting around the attachment point on the DNA. Snapshots from this video are presented in Fig. 4D in the main text. HiLyte 488-labeled tubulin was imaged using 488‑nm illumination. Exposure time: 0.1 s.

Movie S8. Bridging of microtubules by tau-DNA co-condensates

A video of microtubules attached to tau–DNA co-condensates (not shown), bundling together and bridging with one another. HiLyte 488-labeled tubulin was imaged using 488‑nm illumination. Exposure time: 0.1 s.

Movie S9. Changes in high-density DNA surface upon tau and tubulin addition

A video of high-density DNA fluctuations (*top*) and the changes upon the introduction of tau (*middle*) and tubulin (*bottom*). For simplicity, only the DNA (*green*) and tau (*magenta*) fluorescence were merged. The three parts were not recorded simultaneously and were captured from different areas. Snapshots from this video are presented in Fig. 5A–C in the main text. Green: SYTOX Orange-labeled DNA; Magenta: Cy5-labeled tau. Exposure time: 0.1 s.

Movie S10. Microtubules floating over DNA without tau condensate formation

A video of DNA and microtubule fluorescence during a microtubule capture assay. The images were taken over a high-density DNA surface without pre-formed tau condensates. After confirming the absence of microtubules on the surface (0 s), the focus was raised (1 s) to visualize the microtubules flowing over the surface, which failed to interact with the surface due to the absence of pre-formed tau–DNA co-condensates. Snapshots from similar experiments are presented in Fig. 5D in the main text. Top: SYTOX Orange-labeled DNA; Bottom: HiLyte 488-labeled tubulin. Exposure time: 0.1 s.
